# Microstructural and Biomechanical Determinants of Biological Aging

**DOI:** 10.64898/2026.06.16.732769

**Authors:** Pramath Doddaballapur, Jack Di Palo, Dongnan Liu, Liqin Lin, Cristina Cavinato, Abhay B Ramachandra, Xiting Yan, Edward P. Manning

## Abstract

The pulmonary artery undergoes measurable structural and mechanical deterioration with age, but whether these changes can be integrated into a quantitative normative aging prediction model has not been demonstrated. Using two-photon imaging and paired vascular mechanical measurements from C57BL6 mice spanning 6 to 24 months, we developed a multimodal support vector regression (SVR) model integrating collagen fiber orientation, straightness, and hemodynamic mechanical parameters to predict normative age. Fiber orientation was encoded via the von Mises probability density function referenced to the circumferential and axial vessel wall axes providing a principled circular-variable encoding of both mean direction and concentration. The microstructure-only model achieved leave-one-out (LOO) R² = 0.596, Mean Absolute Error (MAE) = 3.43 months. Adding vascular mechanical parameters (PWV) raised a combined LOO R² to 0.834 (MAE = 2.26 months), a 40.1% improvement. Because pulmonary vascular and parenchymal aging are mechanistically coupled, lung mechanics were included as a complementary readout to assess whether airway mechanics contribute independent predictive signal beyond vascular microstructure alone. A sex dimorphism was observed, where females drove the majority of the collagen-based predictive signal (female-only R² = 0.960 vs. male-only R² = 0.658). These results establish a multimodal framework for vascular biological age quantification that integrates structural and mechanical aging signatures.

## Introduction

Cardiovascular aging involves a coordinated remodeling of arterial wall structure and function that unfolds across the lifespan, contributing to increased stiffness, impaired mechanotransduction, and elevated risk of cardiovascular disease [1]. In the pulmonary vasculature, this process is marked by progressive remodeling of the extracellular matrix, particularly collagen, alongside measurable changes in arterial mechanical properties including stiffness, distensibility, and pulse wave velocity [2]. While these structural and mechanical changes have been characterized independently as correlates of vascular aging, prior work has been limited by treating aging as a categorical rather than a continuous variable. Further, we found in previous work that microstructural remodeling contributes to proximal pulmonary arterial stiffening [3][4]. Therefore, it is necessary to evaluate the potential benefit of incorporating microstructure of the proximal pulmonary artery into biomechanical age-related changes. This could establish a unified, quantitative framework that captures biological aging as a continuous, structural-physiologically based process.

Biological clocks predict biological age using biological measurements, providing a quantitative readout of accelerated or attenuated aging [5]. Chronological age measures an individual’s time since birth, whereas biological age quantifies changes in physiological structure and function to reflect age-related declines [6][7]. The majority of established clocks operate on DNA methylation data, but imaging-based and physiological clocks offer complementary advantages, such as mechanistic interpretability, no requirement for molecular assays, and direct grounding in the structural and functional biology of the tissue of interest [3], potentially paving way for noninvasive measures of aging. In our prior longitudinal characterization of the murine cardiopulmonary system, biological age estimated from pulmonary arterial mechanical parameters tracked chronological age across the adult lifespan, demonstrating that physiologically based age clocks are feasible in this system [2]. A multimodal vascular biological age clock integrating 2-photon derived collagen architectural parameters with hemodynamic mechanical measurements represents a new class of aging biomarkers with direct relevance to pulmonary vascular disease.

Vascular mechanical parameters such as arterial distensibility, wall stiffness, and pulse wave velocity, reflect the integrated functional consequence of structural matrix remodeling and provide mechanically interpretable aging correlates [5]. Distensibility measures the fractional change in arterial diameter per unit pressure change and decreases as the wall stiffens with age. Circumferential stiffness quantifies the resistance to radial deformation under physiological loading. PWV is a non-invasive index of arterial stiffness directly tied to the Moens-Korteweg relationship between wall elastic modulus and wave propagation speed [8]. We found that these arterial wall characteristics are sufficient to construct a model of physiologically based biological aging [2]. However, that model was missing micro-structural data such as adventitial collagen orientation, which we previously identified as critical for pulmonary arterial wall stiffening [3][4]. The co-occurrence of collagen architectural remodeling and mechanical stiffening in aging arteries suggests that these two measurement modalities provide complementary information that, when combined, should improve biological age prediction beyond either alone.

We previously found that there are significant increases in circumferential stiffening, collagen reorientation toward the circumferential direction, and vascular cellular senescence in the murine proximal PA between 3 and 24 months [3]. These changes are associated with decreased RV and lung function [3]. Recently, we extended that study by including age as a continuous variable and identifying physiologically based measures of biological, normative aging using principal component analyses of the physiological function [2]. However, that method lacked microstructural contributions. We hypothesized that integrating microstructural and biomechanical properties of components of the cardiopulmonary system with a multimodal model would improve the prediction of normative biological age of the cardiopulmonary system of mice, including sex as a biological variable. To test this hypothesis, we quantified the independent and joint predictive contributions of microstructural collagen parameters and vascular mechanical measurements.

Two-photon microscopy provides quantitative access to fibrillar collagen architecture without tissue disruption or chemical labeling [9]. Two aspects of collagen organization are particularly informative for vascular aging: the preferred orientation of collagen fibers relative to the vessel wall axes, and the degree to which individual fibers are geometrically taut versus crimped. With age, collagen fibers in the pulmonary arterial wall progressively reorient toward the circumferential direction under chronic pressure loading, while simultaneously losing their characteristic crimp and becoming increasingly straight. These structural changes reflect the cumulative mechanical and biochemical remodeling of the extracellular matrix and serve as interpretable histological correlates of vascular age.

We report that integrating vascular microstructure with biomechanical parameters substantially improves biological age prediction, with lung mechanics providing an independent contribution. We further characterize a sex dimorphism in which the collagen aging signal is substantially stronger in females than in males.

## Methods

### Animals and tissue preparation

We analyzed cardiac and pulmonary arterial data acquired during a normative aging mouse study reported elsewhere [2]. Briefly, C57BL6 mice from Jackson Laboratories or NIA Aging colony were maintained under standard housing at Yale University and protocols were approved by Yale University Institutional Animal Care and Use Committee. Age groups included 3, 6, 12, 18, 24, and 27 months. Three-month animals were pooled with 6-month animals due to no established vascular aging phenotype distinguishing 3 month from 6 month in this strain; 27-month animals were pooled with 24 month animals to maintain a 6–24-month prediction range and handle edge-case continuity errors.

### Two-photon imaging and collagen orientation extraction

The RPA was dissected, tested, and imaged as described in De Man et al. [3]. Briefly, the right main pulmonary artery from the bifurcation of the main pulmonary artery to the first branch of the right pulmonary artery was excised and canulated on custom glass micropipettes. The artery and any branches include the ligamentum arteriosum were ligated with suture, thus securing the artery onto the glass cannulas and preventing leaks. After mechanical testing, the arteries were cannulated onto metal overlapping blunt needles and ligated at each end, thus allowing pressurization of the artery while also controlling its stretch.

Representative pulmonary-artery regions were imaged under *in vivo*–relevant stretches and pressures (matching passive mechanical testing). Multiphoton imaging used a Titanium-sapphire laser (Chameleon Vision II, Coherent) on a TriMScope microscope (LaVision Biotec) tuned to 820 nm with a 20× water-immersion objective (NA 0.95). Backward scattering second harmonic vgeneration signal (SHG) from fibrillar collagen was collected at 390–425 nm. Volumes (500 × 500 µm field of view; ∼0.05 mm³) were acquired at 0.48 µm/pixel in-plane resolution with 1 µm z-steps. Three-channel 3D stacks were post-processed in MATLAB R2024b and ImageJ 1.53a by fitting a circle to the mid-wall profile and transforming circumferential–radial slices from polar to Cartesian coordinates to enable layer-specific collagen alignment and cell volume-density analyses [10]. Images were then cropped to a 1020 × 1010-pixel region of interest to remove edge artifacts [11].

Two-photon imaging dataset comprise 81 samples (n = 81). Paired vascular mechanical measurements were available for subsets of the same animals (n = 29–41 depending on mechanical parameter; see below).

Collagen fiber orientation was quantified using the OrientationJ Distribution plugin of Fiji from the SHG channel. For each image slice, a pixel-wise local orientation map was computed using the structure tensor method and exported the orientation frequency histogram (angle in degrees vs. normalized pixel count). The following OrientationJ parameters were used: structure tensor window σ = 2.0 pixels; gradient method = cubic spline (gradient = 4); minimum coherency threshold = 20%; minimum energy threshold = 5%. The von Mises distribution parameters θ (mean orientation, degrees) and κ (concentration, dimensionless) were estimated by fitting the von Mises probability model to each exported histogram using maximum likelihood estimation in Python (scipy.stats.vonmises).

### Von Mises Feature Construction

Raw θ and κ were not used directly as model inputs. Instead, vm_circ and vm_axial were derived as the probability mass of the fitted von Mises distribution integrated over the circumferential and axial angular windows, respectively:

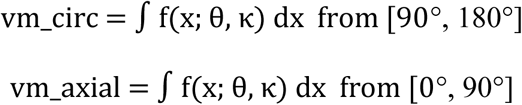

where *f*(*x*; θ, *x*) = 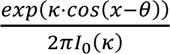 is the von Mises PDF, θ is the mean fiber orientation angle, κ is the concentration parameter (inversely related to fiber dispersion), and I₀(κ) is the modified Bessel function of the first kind of order zero. This formulation captures the full fiber orientation distribution within each angular window rather than a point evaluation and naturally accommodates fibers at intermediate angles by apportioning their contribution proportionally between the two windows. Because the von Mises distribution integrates to unity over the full circle, vm_circ and vm_axial sum to 1 and are therefore negatively coupled: increasing circumferential alignment necessarily reduces axial density. The von Mises distribution is the natural model for angular data; it is periodic, handles the 0°/360° wrap-around correctly, and its concentration parameter κ directly quantifies the degree of fiber alignment.

### Collagen straightness quantification

Collagen straightness was defined as the ratio of end-to-end Euclidean chord length to total curvilinear fiber contour length (range 0–1; 1 = perfectly straight). Two independent measurements (automated and manual) were made per image and averaged to produce the final value. For automated measurements, CT-FIRE software [12] extracted individual fiber centerline coordinates from each image and computed per-fiber straightness; mean straightness across all CT-FIRE- detected fibers was taken as the automated estimate. For manual measurements 10 individual fibers per image were selected and manually traced by the analyst; the chord-to-contour ratio was computed for each and averaged. The final straightness value for each sample is the mean of the CT-FIRE mean and the manual mean. This dual-method approach reduces dependence on automated fiber detection in images with variable signal quality. Together, straightness and Von Mises orientation fully characterize the state at the two mechanically relevant vessel wall axes. The three collagen features used in all models were: vm_circ, vm_axial, and collagen straightness.

### Vascular mechanical measurements

PA structural and mechanical parameters were measured by biaxial tensile testing as previously described [2][3]. Briefly, excised RPA segments were mounted on a biaxial testing apparatus and pressurized over a range of physiological pressures at fixed axial stretch. The following parameters were extracted from the structural and material relationships: PA loaded outer diameter (μm), PA loaded wall thickness (μm), PA axial and circumferential stretches (dimensionless), PA elastic stored energy (kPa), PA axial and circumferential stresses (kPa), arterial distensibility (mmHg⁻¹), PA circumferential stiffness (MPa), PA axial stiffness (MPa), and pulse wave velocity (PWV, m/s), computed from the Bramwell-Hill equation applied to the measured compliance. Images were obtained and biomechanical properties calculated at specimen-specific pressures and in vivo axial stretch.

Not all parameters were available for every animal; the present analysis used a subset of animals with complete orientation data and the specific mechanical parameters included in each model configuration. For the primary multimodal model (2-photon + distensibility + PA circumferential stiffness + PWV), n = 36 animals had all five features.

### Lung mechanics measurements

Lung mechanics were measured by forced oscillation technique (flexiVent, SCIREQ) [13]. Animals were anaesthetized, tracheotomized, and mechanically ventilated at 150 breaths/min, 10 mL/kg tidal volume, 3 cmH₂O PEEP. The following parameters were extracted from the constant-phase model fit to the input impedance spectrum: airway resistance Rn (cmH₂O·s/mL), tissue damping G (cmH₂O/mL), tissue elastance H (cmH₂O/mL), and hysteresivity η = G/H (dimensionless). Static compliance Cst and dynamic compliance Ers were also recorded. Lung mechanics data were available for a partially overlapping subset of animals relative to the 2-photon dataset (n = 29 with complete collagen and lung resistance data in the 6–24 month range). Critically, lung resistance and the PA mechanical parameters (distensibility, stiffness, PWV) were measured in entirely separate animal cohorts within the same age groups, meaning their samples do not overlap, precluding any combined 2-photon + lung + PA mechanical model.

### Preprocessing pipeline

Within each LOO fold, training samples were preprocessed to satisfy the assumptions of kernel-based learning. First, samples were winsorized at the 5th–95th percentile to limit the influence of outliers, which can disproportionately distort the RBF kernel’s distance calculations. Second, a Yeo-Johnson power transformation was applied per feature to reduce skewness and approximate normality, improving the interpretability of kernel distances in the transformed feature space. Third, features were standard scaled to zero mean and unit variance to ensure that no single feature dominated the RBF kernel by virtue of its scale rather than its biological relevance. The test sample was transformed using training-fold parameters only, strictly preventing data leakage.

### Support vector regression: full mathematical formulation

Support vector regression (SVR) was selected to capture the relationship between collagen microstructure, vascular mechanics, and chronological age. Biological aging processes are governed by nonlinear interacting mechanisms, such as collagen crosslinking, fiber reorientation, and vascular stiffening that do not accumulate linearly with time, making linear regression an insufficient model. SVR addresses this through a kernel, which implicitly maps input features into a high-dimensional space where nonlinear relationships become linearly separable, without requiring explicit specification of the transformation. Additionally, SVR’s margin-based loss function (the ε-insensitive tube) provides robustness to small residuals and outliers, which is advantageous given the biological variability inherent in a small animal cohort.

SVR minimizes the regularized primal objective:

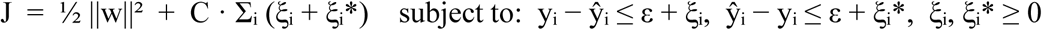

where ||w||² penalizes model complexity (the norm of the weight vector in kernel space); C = 200 controls how strongly constraint violations are penalized; ξᵢ and ξᵢ* are per-sample slack variables measuring excess error above and below the ε-tube respectively; and L_ε_(yᵢ − ŷᵢ) = max(0, |yᵢ − ŷᵢ| − ε) is the epsilon-insensitive loss with ε = 1.5 months. The regularization parameter C was selected via grid search (C ∈ {1, 10, 50, 100, 200, 500, 1000}) under LOOCV, with C = 200 minimizing cross-validated MAE across all feature configurations.The Lagrangian of the primal is formed by associating dual variables αᵢ ≥ 0, αᵢ* ≥ 0, and μᵢ, μᵢ* ≥ 0 with each primal constraint. Stationarity with respect to w and b (the intercept) yields the KKT conditions: w = Σᵢ(αᵢ − αᵢ*)ϕ(xᵢ) and Σᵢ(αᵢ − αᵢ*) = 0. Complementary slackness conditions require that: αᵢ(ε + ξᵢ − yᵢ + ŷᵢ) = 0 and αᵢ*(ε + ξᵢ* + yᵢ − ŷᵢ) = 0, meaning that only samples outside the ε-tube (ξᵢ > 0 or ξᵢ* > 0) have non- zero dual variables and become support vectors. Additionally, μᵢ · ξᵢ = 0, so samples with ξᵢ > 0 have μᵢ = 0 and therefore their αᵢ can equal C, meaning that these are fully saturated boundary support vectors, the hardest-to-fit training samples.

Substituting the stationarity condition into the Lagrangian and eliminating the primal variables provides the dual objective:

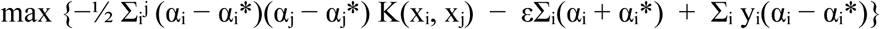

subject to Σᵢ(αᵢ − αᵢ*) = 0 and 0 ≤ αᵢ, αᵢ* ≤ C. The kernel function K(xᵢ, xⱼ) = 〈ϕ(xᵢ), ϕ(xⱼ)〉 replaces explicit computation of the high-dimensional feature map ϕ. The RBF kernel K(xᵢ, xⱼ) = exp(−γ||xᵢ − xⱼ||²) with γ = 0.05 corresponds to an infinite-dimensional feature space containing products of all degrees, enabling nonlinear regression without explicit feature expansion. The dual is a quadratic program with O(n²) terms, solved efficiently by sequential minimal optimization (SMO) as implemented in scikit-learn[14]. The weight norm in kernel space is: ||w||² = Σᵢʲ (αᵢ − αᵢ*)(αⱼ − αⱼ*) K(xᵢ, xⱼ), quantifying model complexity as the total interaction strength among support vectors weighted by their dual coefficients. The prediction equation is:

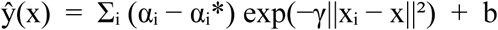

For the 2-photon-only model: 50 of 86 training samples became support vectors; 36 reached the ±C boundary; bias b = 6.46 months. Positive dual coefficients (true age above the tube) pull predictions toward older ages about that support vector; negative coefficients pull toward younger ages. The prediction is therefore a weighted sum of Gaussian similarity scores between the test point and each support vector, with weights reflecting both how well each support vector was fitted and the polarity of the fitting error. Hyperparameters C = 200, ε = 1.5, γ = 0.05 were selected by grid search maximizing LOO R² on the full 2-photon-only dataset.

### Model configurations evaluated

Four model configurations were evaluated and are reported as primary results: (1) 2-photon only: vm_circ + vm_axial + collagen straightness, n = 81; (2) 2-photon + distensibility: n = 45; (3) 2-photon + distensibility + PA circumferential stiffness: n = 45; (4) 2-photon + distensibility + PA circumferential stiffness + PWV (primary multimodal model): n = 36. Two additional configurations are reported as exploratory secondary findings: (5) 2-photon + lung resistance: n = 29; (6) 2-photon + lung resistance + hysteresis: n = 29. The lung mechanics and PA mechanical cohorts have zero sample overlap and cannot be combined. All configurations used identical hyperparameters and preprocessing.

### Cross-validation and statistical analysis

LOOCV was the primary validation method. With a dataset of n = 81 samples distributed across four age groups and two sexes, the choice of cross-validation strategy has a meaningful impact on both the reliability of the performance estimate and the amount of data available for training at each fold. Standard k-fold cross-validation partitions the dataset into k equal groups, trains on k−1 groups, and tests on the remaining group. At k = 5, each training fold would contain only 69 samples, and at k = 10, only 77. Given that the dataset is already modest in size and further stratified across age and sex, each reduction in training set size risks destabilizing the model, particularly at the age group boundaries where sample counts are lowest. LOOCV addresses this directly by training on n−1 = 85 samples at every fold and testing on a single held-out observation, maximizing the training set size at every step. This comes at a computational cost (86 separate model fits rather than 5 or 10), but is feasible for SVR at this sample size, and is well-justified given the high quality and consistency of the training data, which was acquired under a standardized single-laboratory protocol.

Furthermore, LOOCV produces an approximately unbiased estimate of generalization error for small datasets, whereas k-fold estimates carry higher variance when k is small relative to n. For the sex-stratified analyses, where subsets shrink to n = 18 (males) and n = 18 (females), the case for LOOCV is even stronger: To maximize the amount of training data available at each fold, we used leave-one-out cross-validation (LOOCV) rather than k-fold. With only 32 samples, a 5-fold approach would train on just 25 samples per fold, too few to reliably fit a nonlinear model. In LOOCV, the model trains on all but one sample and is tested on the held-out sample, repeated for every sample in the dataset. To prevent data leakage, all preprocessing steps were fit exclusively on the training samples at each fold and then applied to the test sample. Spearman correlations assessed individual parameter–age relationships. Mann-Whitney U tests assessed sex differences at each age group. All analyses used Python 3.12 with scikit-learn [14], scipy [15], and numpy [16]. Significance threshold α = 0.05, two-tailed. Model significance was assessed by a permutation test: age labels were randomly shuffled 1,000 times, the full SVR-RBF pipeline (winsorization, Yeo-Johnson, standardization, LOO prediction) was refitted on each permuted dataset, and the proportion of permuted LOO R² values meeting or exceeding the observed LOO R² was taken as the empirical p-value. Age-group differences in collagen parameters were assessed by Kruskal-Wallis test followed by Dunn’s post-hoc test with Bonferroni correction, stratified by sex (Supplementary Table 1).

### Validation

To validate our model, we predicted the chronological age of 10 mice spanning the full age range of the training dataset (3–28 months, Table 2). Input features including hemodynamic parameters (distensibility, PWV, PA circumferential stiffness) and 2-photon collagen morphology metrics (θ, κ) were extracted and submitted to the trained SVR model without investigator knowledge of group assignment. True chronological ages were revealed only after all predictions were recorded.

**Table 1.** Age-related changes in murine cardiopulmonary system are best fit by analyzing variance in microstructural properties (collagen orientation and straightness) and proximal pulmonary artery distensibility and wall stiffness. Model performance across 2-photon and multimodal configurations (6–24mo) are shown. LOO = leave one out. MAE = mean average error (in months). Green: 2-photon-only baseline. Blue: primary multimodal model. Lung mechanics and PA mechanical cohorts have zero overlap and cannot be combined.

| Model features | n | LOO R<sup>2</sup> | MAE (mo) | vs. 2-photon baseline |
| --- | --- | --- | --- | --- |
| 2-photon only (vm_circ + vm_axial + Collagen) | 81 | 0.596 | 3.43 | Reference |
| 2-photon + Distensibility | 45 | 0.740 | 3.08 | +0.145 |
| 2-photon + Distensibility + PA Circ. Stiffness | 45 | 0.818 | 2.53 | +0.202 |
| **2-photon + Distensibility + PA Circ. Stiffness + PWV [Primary multimodal]** | 36 | 0.834 | 2.26 | +0.239 |
| 2-photon + Lung Resistance [Exploratory] | 29 | 0.854 | 2.22 | +0.258 |
| 2-photon + Lung Resistance + Hysteresis [Exploratory] | 29 | 0.797 | 2.60 | +0.222 |

**Table 2.** Blind validation cohort: SVR-predicted vs. chronological age.

| Mouse ID | Group | Actual Age (mo) | Predicted Age (mo) | Error (mo) |
| --- | --- | --- | --- | --- |
| A | 3 mo, Male | 3 | 4.03 | +1.03 |
| B | 3 mo, Male | 3 | 4.02 | +1.02 |
| C | 5 mo, Female | 5 | 7.30 | +2.30 |
| D | 5 mo, Female | 5 | 7.40 | +2.40 |
| J | 22 mo, Male | 22 | 20.14 | −1.86 |
| K | 22 mo, Male | 22 | 16.69 | −5.31 |
| L | 25 mo, Female | 25 | 24.10 | −0.90 |
| M | 25 mo, Female | 25 | 22.64 | −2.36 |
| N | 6 mo, Female | 6 | 8.12 | +2.12 |
| P | 28 mo, Male | 28 | 24.90 | −3.10 |

*Mouse L (best prediction, error −0.90 mo) and Mouse K (outlier, error −5.31 mo)*

## Results

We found that age-related changes in murine proximal pulmonary arteries are best fit by analyzing variance in microstructural properties (collagen orientation and straightness) and proximal pulmonary artery distensibility and wall stiffness, as shown in Table 1. We also found that age-related changes in pulmonary artery microstructure, distensibility, and wall stiffness significantly differ between age groups and sex, as shown in Supplemental Table S1 and Supplemental Figure S1.

### Collagen architectural parameters track age monotonically

Across the full n = 81 dataset, κ declined monotonically with age (Spearman ρ = −0.518, p < 0.0001), collagen straightness increased monotonically (ρ = +0.581, p < 0.0001), and mean fiber orientation angle θ increased monotonically (ρ = +0.419, p < 0.001;Figure 1), reflecting progressive reorientation of collagen fibers toward the circumferential vessel wall direction with advancing age. Kruskal-Wallis tests confirmed significant age-group variation in all three parameters in females (κ: H = 27.89, p < 0.001; straightness: H = 24.61, p < 0.001; θ: H = 13.22, p = 0.004), while male κ did not reach significance (H = 7.07, p = 0.070), consistent with the attenuated collagen aging trajectory observed in male animals. Dunn post-hoc pairwise comparisons (Bonferroni correction) confirmed significant female κ decline between the 6–12 mo and 24 mo groups (p < 0.001 for both), with no significant pairwise differences in male κ after correction (Supplemental Table S1, Supplemental Figure S1). Within individual age groups, κ, θ, and collagen straightness were generally non-significantly correlated with one another (p > 0.10 for most pairwise comparisons), indicating that they capture largely independent aspects of collagen structural aging, except for a significant κ–straightness association at the 18-month timepoint (p = 0.006, n = 13). Collagen dispersion from 6 to 24 months increases significantly in female mice, but not males. Female κ fell from 8.61 ± 2.25 at 6 months to 2.20 ± 2.02 at 24 months, a 74% decline; male κ declined only 14% over the same interval (5.83 to 4.99). Mean θ increased in both sexes with age, consistent with the circumferential fiber reorientation documented in De Man et al., but the increase was more pronounced and consistent in females. The circumferential von Mises feature vm_circ integrates both of these trends simultaneously, where a fiber population that is both more circumferentially oriented (higher θ toward 90°) and more concentrated (higher κ) produces a disproportionately large vm_circ value, which explains the 90-fold increase in females (0.0012 to 0.104) compared to the 4.5-fold increase in males (0.0038 to 0.017). The vm_circ trajectory thus amplifies the combined signal of θ and κ aging, making it a more sensitive aging readout than either parameter alone.

**Figure 1.**
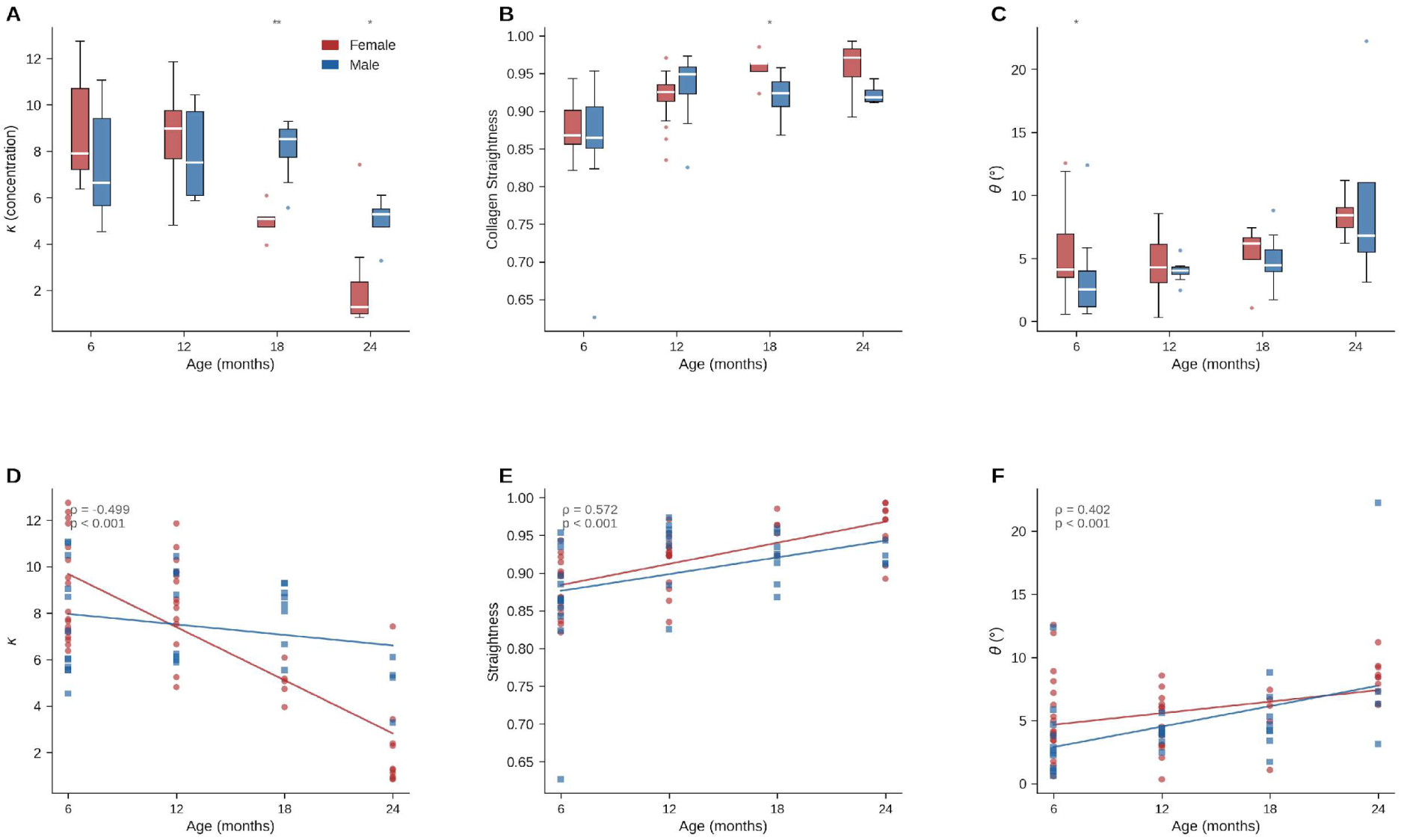
Collagen orientation, anisotropy (misalignment), and straightness in the walls of the proximal pulmonary artery are key age-related microstructural changes in the cardiopulmonary system of mice. Collagen architectural parameters by age group and sex. (A) κ, (B) collagen straightness, and (C) θ by age group, stratified by sex (blue, male; red, female). κ declines monotonically (Spearman ρ = −0.518, p < 0.001), straightness increases (ρ = +0.581, p < 0.001), and θ increases (ρ = +0.419, p < 0.001). (D–F) Spearman correlation scatter plots vs. chronological age. Mann-Whitney U: *p < 0.05, **p < 0.01, ***p < 0.001. n = 81.

### Microstructural changes of the pulmonary artery correlates with age

The three-feature 2-photon model (vm_circ + vm_axial + collagen straightness) achieved LOO R² = 0.596, MAE = 3.43 months (n = 81; Figure 2, Table 1), representing an 8.1% improvement over linear regression on identical features (LOO R² = 0.551). Model significance was confirmed by permutation test: zero of 1,000 label-permuted models achieved LOO R² ≥ 0.596 (p < 0.001), confirming a nonlinear age–morphometry relationship. The optimal model used three features, since incorporating the remaining five parameters from the 2-photon which were: adventitia thickness, media thickness, adventitial cell count, SMC count, and endothelial cell count degraded performance to R² = 0.417, indicating that additional structural features introduced noise rather than predictive signal.

**Figure 2.**
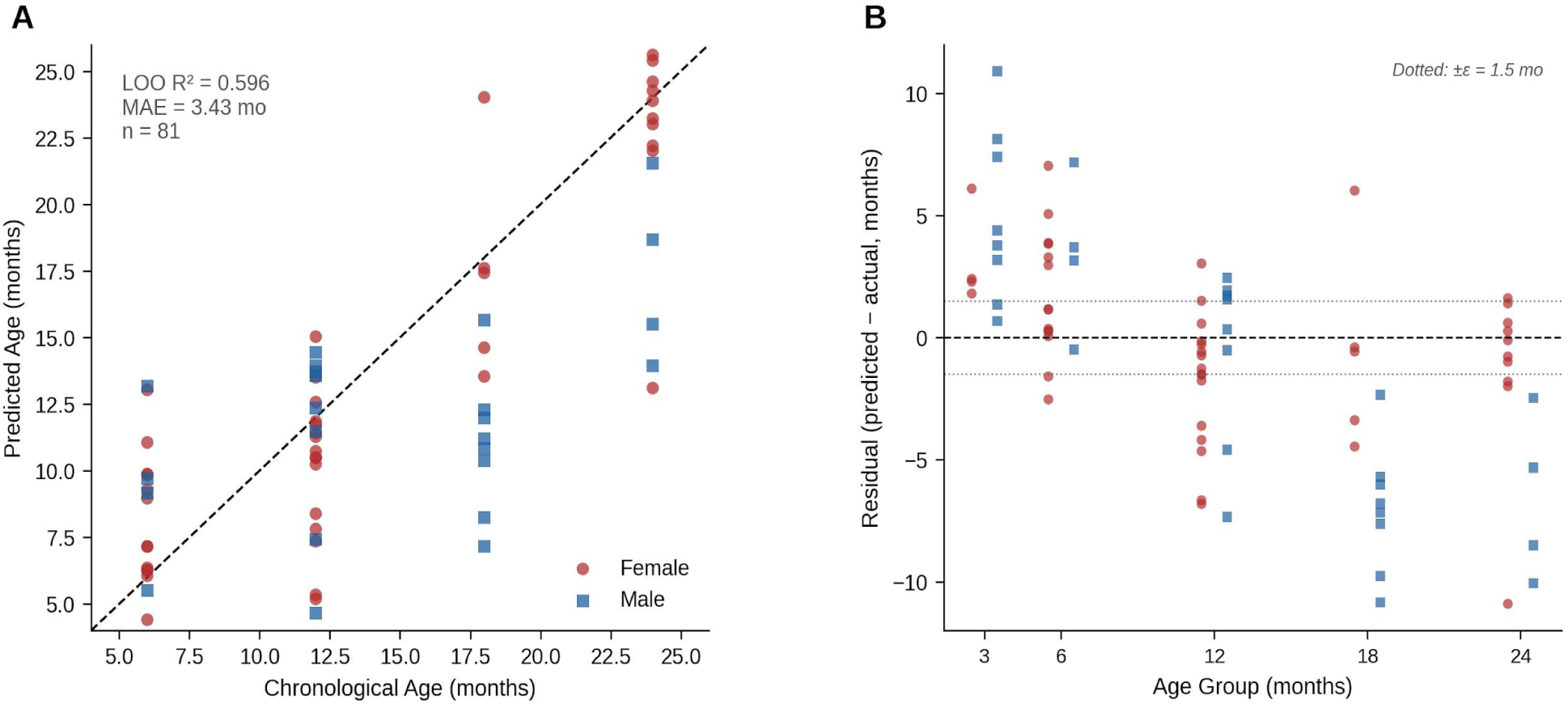
A model trained on microstructure of the proximal pulmonary artery can predict chronological age. 2-photon-only SVR-RBF model: LOO cross-validation (n = 81). (A) Predicted vs. chronological age (LOO R² = 0.596, MAE = 3.43 months); dashed line, identity; red circles, female; blue squares, male. (B) Residuals by age group; dotted lines, ±ε = 1.5-month tolerance.

### Multimodal modeling of microstructural and mechanical properties of the pulmonary artery provide a better model of normative aging

To improve our model of normative aging of the proximal pulmonary artery, we included mechanical properties of the pulmonary artery into a multimodal model. We found that adding three PA mechanical parameters to the collagen features raised LOO R² from 0.596 to 0.834 (MAE = 2.26 months, n = 36), a 40.1% absolute improvement (Figure 3A, Table 1). The stepwise improvement is informative: distensibility alone increased R² to 0.740 (n = 45); distensibility + PA circumferential stiffness reached 0.818 (n = 45); the full combination with PWV reached 0.834 (n = 36). The improvement from distensibility reflects the direct mechanical consequence of collagen matrix aging with progressive loss of vascular compliance as fibers uncrimp, cross-link, and bear load, providing the model with a functional correlate of the structural remodeling captured by the 2-photon features. PA circumferential stiffness adds material information about the resistance to radial deformation under pressure, reflecting wall stiffening associated with extracellular matrix remodeling. PWV, as an integrated measure of wall elasticity across the vessel length, captures a globalized mechanical aging signature not fully described by local structural measurements alone.

**Figure 3.**
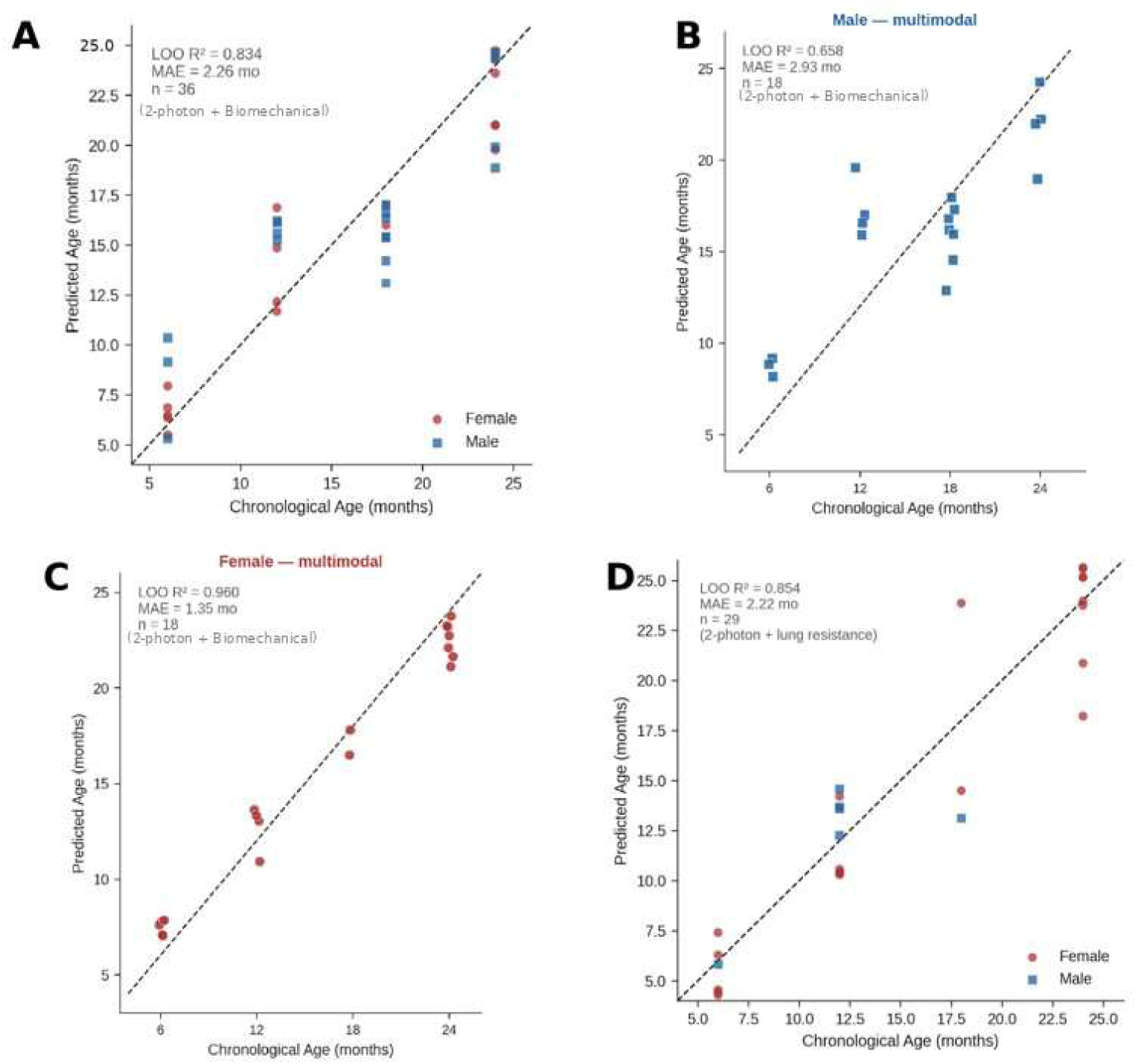
Multimodal SVR models predict biological age of the murine proximal pulmonary artery across sexes and organ compartments. LOO cross-validation scatter plots of SVR-predicted vs. chronological age for each model configuration. (A) Primary multimodal model combining 2-photon collagen features, distensibility, PA circumferential stiffness, and PWV (n = 36, LOO R² = 0.834, MAE = 2.26 months). (B) Male-only stratified analysis of the primary multimodal model (n = 18, LOO R² = 0.658, MAE = 2.93 months). (C) Female-only stratified analysis (n = 18, LOO R² = 0.960, MAE = 1.35 months) (D) Exploratory model combining 2-photon features with lung airway resistance (n = 29, LOO R² = 0.854, MAE = 2.22 months), reflecting shared ECM aging biology between the pulmonary vasculature and airways

We then combined lung mechanics as an additional age-predictive signal, reflecting shared pulmonary aging biology. RV functional parameters were excluded due to their collinearity with PA mechanical features already present in the model. As a downstream hemodynamic consequence of pulmonary vascular stiffening, RV remodeling indices carry redundant predictive information that risks degrading SVR generalization in a *small-n* setting.

### Sex dimorphism in collagen-based age prediction

We sought to determine the role of sex as a biological variable in the aging process of the cardiopulmonary system of mice. We found that female-only LOOCV (n = 18) on the combined 2-photon + mechanistic model achieved LOO R² = 0.960 (MAE = 1.35 months). Male-only LOOCV (n = 18) achieved LOO R² = 0.658 (MAE = 2.93 months; Figure 3B). The female κ trajectory shows a nonlinear post-18-month acceleration, producing high feature contrast across the age range (Figure 3C). Male κ is comparatively flat through 18 months, reducing signal.

### Lung mechanics as a secondary age-predictive signal

We further improved our model by including lung mechanical properties of the respective age groups. Adding lung airway resistance (Rn) to the 2-photon features raised LOO R² from 0.596 to 0.854 (MAE = 2.22 months, n = 29), the largest single-feature gain observed in this study (Figure 3D). Lung G (tissue damping, R² = 0.758), lung H (tissue elastance, R² = 0.753), hysteresis (R² = 0.734), and Ers (respiratory system elastance, R² = 0.759) all showed similar gains. Adding lung resistance and hysteresis together gave R² = 0.797; lung resistance alone was the optimal parsimonious choice. However, lung resistance and the PA mechanical parameters derive from entirely separate animal cohorts within the same age groups, meaning they share no specific animal overlap, but only age continuity.

### PCHIP parameter trajectories

Piecewise Cubic Hermite Interpolating Polynomial (PCHIP) spline trajectories through per-sex age-group means confirm nonlinear aging kinetics (Figure 4). Both sexes show a monotonic decline in κ from 6 to 24 months, with females declining more steeply. Collagen straightness rises sharply from 6 to 12 months and plateaus thereafter. Fiber orientation angle θ increases progressively with age in females, with greater sex divergence at later time points. PA distensibility remains stable across mid-life with a male-specific rise at 24 months. Together, these trajectories confirm that all four parameters change directionally and systematically with age, validating their biological relevance as aging correlates.

**Figure 4.**
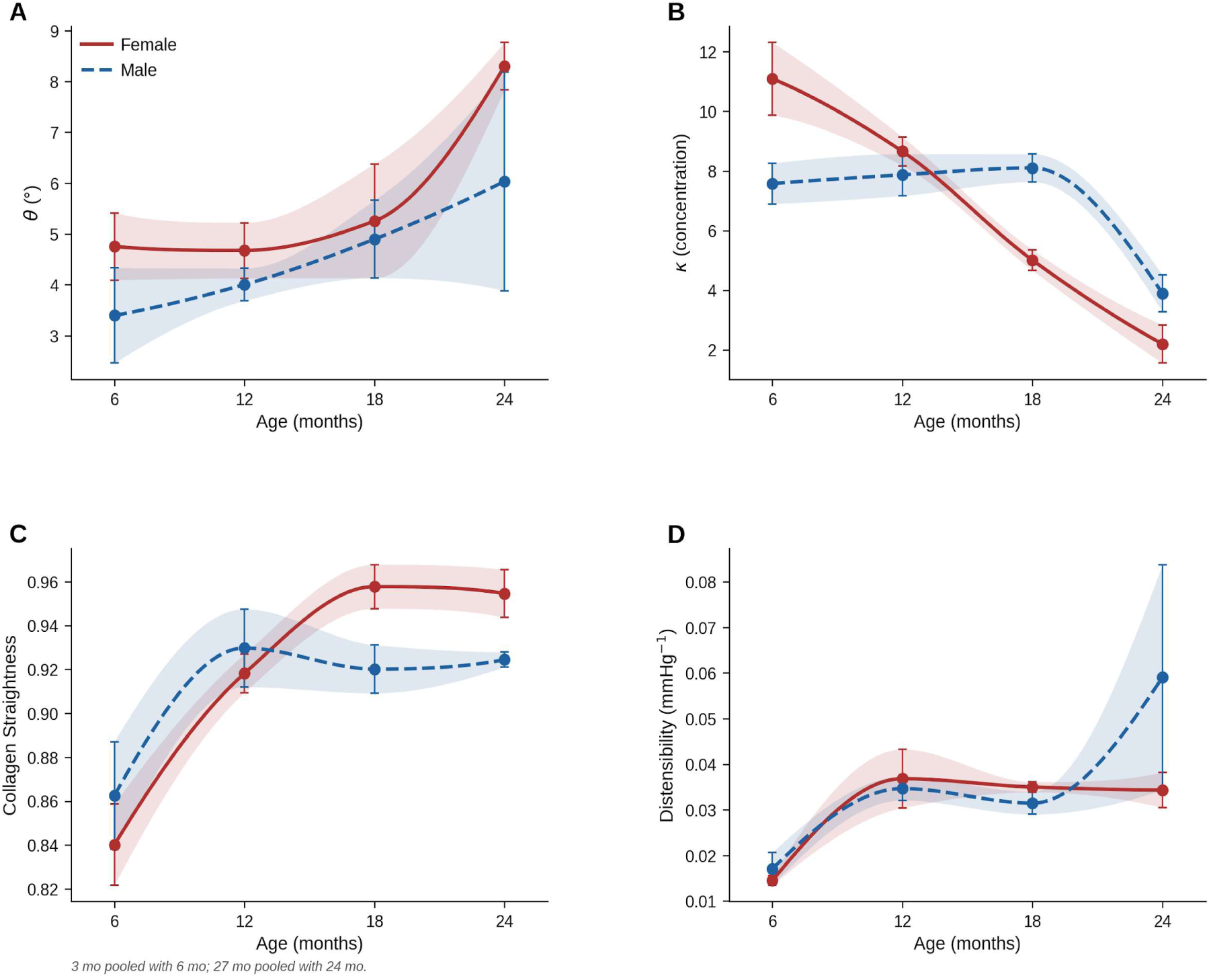
Predicted cardiopulmonary-specific variable as a function of age. Age-related trajectories of collagen architectural parameters and PA distensibility by sex. PCHIP splines connect per-sex age-group means ±SEM (shaded bands). 3 mo animals are pooled with the 6 mo group; 27 mo animals are pooled with the 24 mo group. (A) Fiber orientation angle θ, which increases progressively with age in females and shows greater variability in males. (B) Von Mises concentration parameter κ, reflecting collagen anisotropy. (C) Collagen straightness, which rises rapidly from 6 to 12 months and plateaus thereafter in both sexes (D) PA distensibility (mmHg⁻¹), which increases modestly from 6 to 12 months then stabilizes.

### Permutation feature importance identifies PWV and PA circumferential stiffness as dominant predictors

To assess the relative contribution of each feature to the primary multimodal model, we performed permutation feature importance analysis (80 permutations per feature; Figure 5). Among the mechanical parameters, PWV and PA circumferential stiffness produced the largest mean R² decreases upon permutation, indicating that bulk arterial stiffness and wave propagation speed carry much of the mechanical age signal. This is consistent with the mechanistic role of these parameters as integrated readouts of wall composition and load-bearing behavior across the full vessel length, properties that change directionally and substantially over the 6–24 month aging window. Distensibility, while the first mechanical parameter added in the stepwise model and responsible for the largest single-step R² gain, contributed less to permutation importance than PWV and circumferential stiffness, likely reflecting its higher collinearity with the 2-photon collagen features. Among the microstructural parameters, vm_circ and collagen straightness were the strongest 2-photon contributors, consistent with their monotonic age trajectories across the full dataset. vm_axial showed lower importance, as expected given its negative coupling with vm_circ under the von Mises normalization constraint. Together, the feature importance profile confirms that the multimodal model draws on genuinely complementary information from both modalities, with no single feature dominating, and supports the biological interpretability of the aging signal captured by the SVR framework.

**Figure 5.**
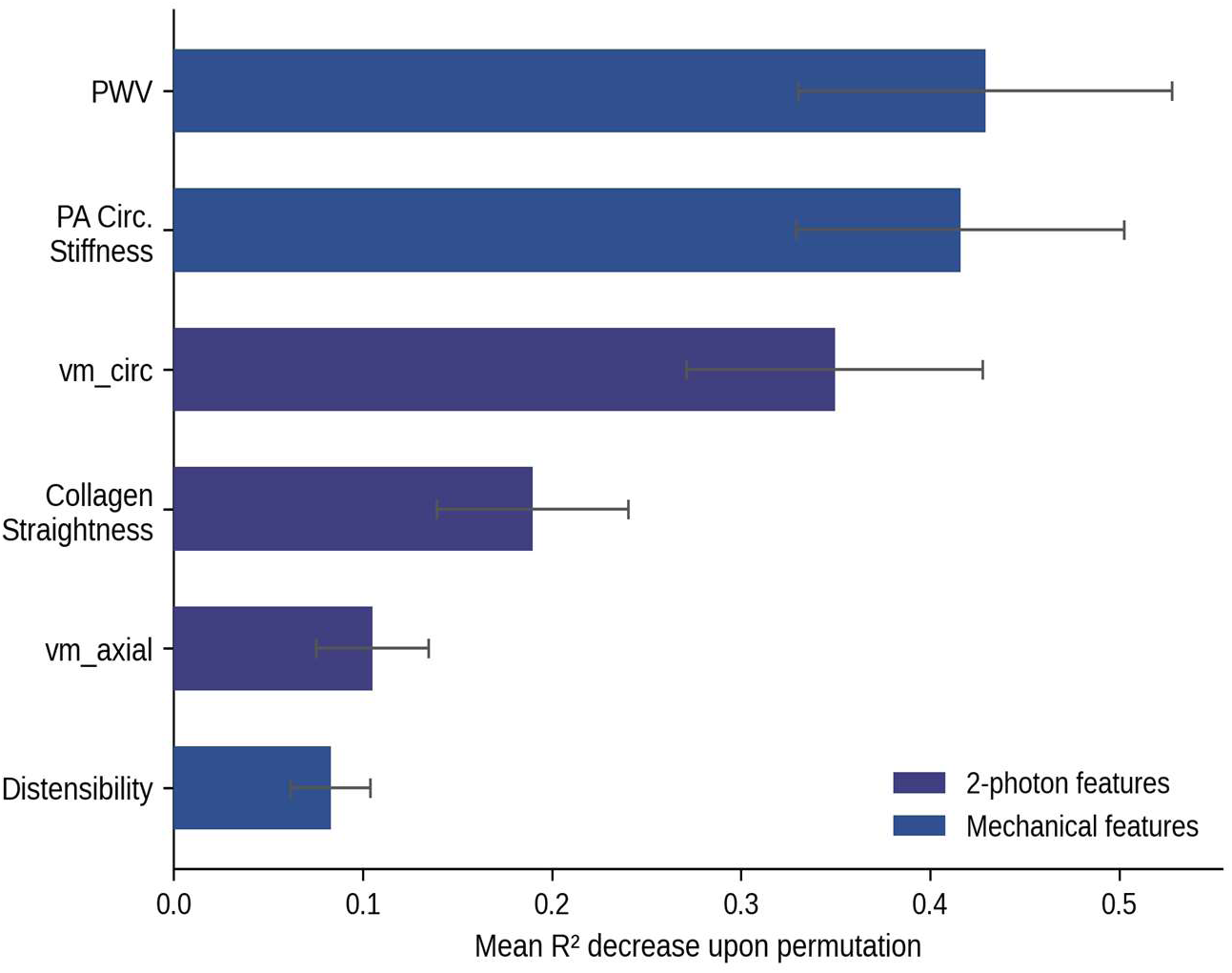
Correlations of structural and function components of the proximal pulmonary artery on age-related changes. Permutation feature importance — primary multimodal model (n = 29, 80 permutations). Mean R² decrease upon permutation (±1 SD). PWV and PA circumferential stiffness are the dominant mechanical features; vm_circ and collagen straightness are the strongest 2-photon features. Error bars, ±1 SD across 80 permutations.

### Predictive Accuracy

The model demonstrated strong predictive accuracy on the blind cohort, achieving an R² of 0.937 and a mean absolute error (MAE) of 2.24 months, comparing favorably to cross-validated training performance (R² = 0.834, MAE = 2.26 months) and suggesting the model generalizes well to unseen animals (Figure 6). Of the 10 blinded predictions, 8 (80%) fell within ±3.0 months of the true chronological age. Based on the fitted residual distribution (μ = −0.47 mo, σ = 2.65 mo; Shapiro-Wilk p = 0.303), the estimated probability of any given prediction falling within ±3.0 months is 75.1%. Taken together, these results support the robustness of the multimodal SVR framework for pulmonary vascular biological age estimation in mouse cohorts.

**Figure 6.**
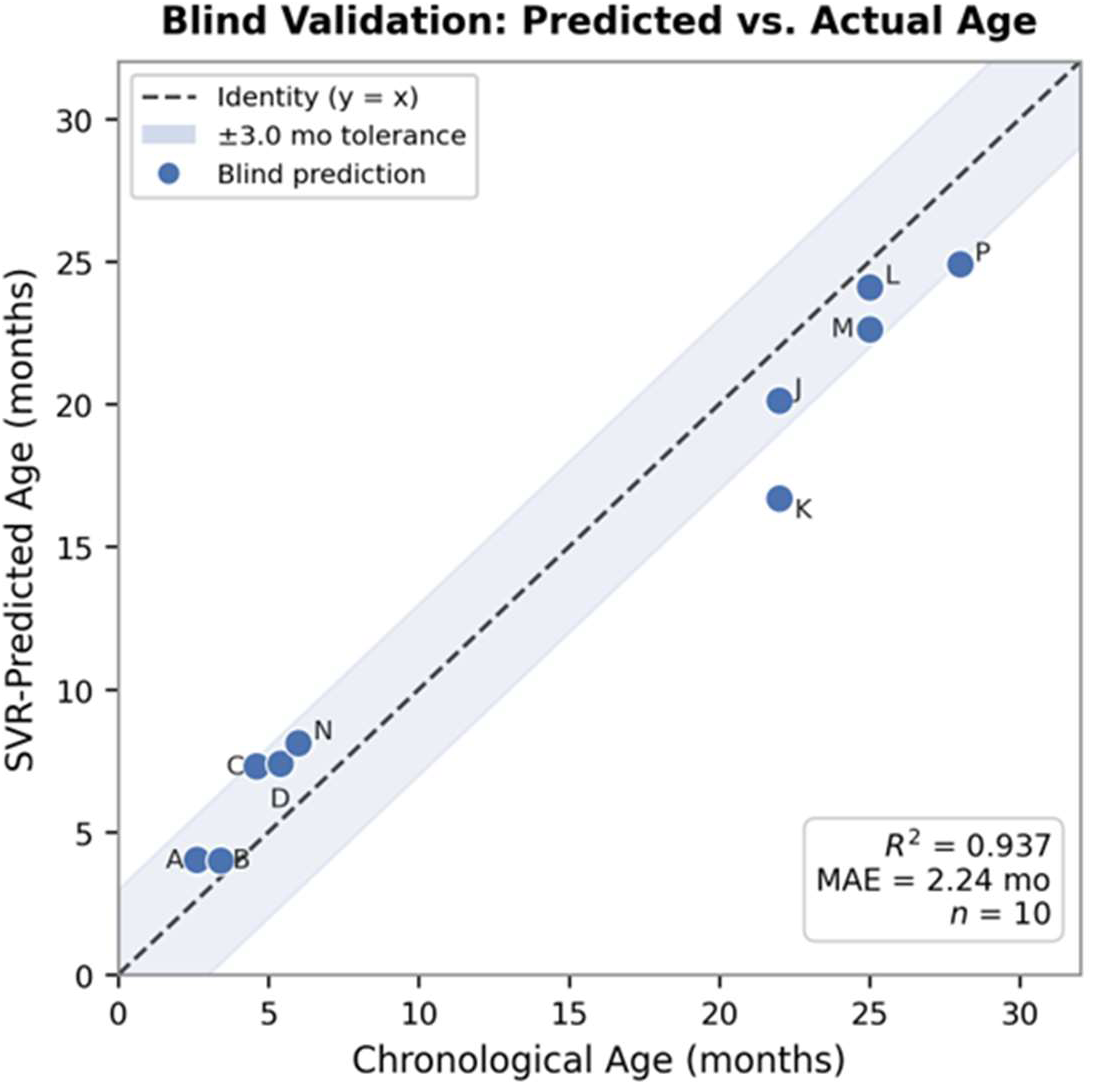
Correlation between SVR-predicted and chronological age in a blinded validation cohort (n = 10). Dashed line indicates the identity (y = x). Shaded region denotes ±3.0-month tolerance. R² = 0.937, MAE = 2.24 months.

## Discussion

We hypothesized that integrating microstructural and biomechanical properties of components of the cardiopulmonary system with a multimodal system model can predict age of the cardiopulmonary system of mice. We found that the primary multimodal model which combines von Mises orientation features, collagen straightness, distensibility, PA circumferential stiffness, and pulse wave velocity, achieved LOO R² = 0.834, reliably predicting the age of normative aging of the cardiopulmonary system of mice. The improvement reflects the complementary nature of structural and biomechanical aging information: collagen features capture the microstructural organizational state of the fiber network, while biomechanical parameters of the PA capture the functional consequence of normative age-related remodeling. This study presents the first multimodal pulmonary vascular aging model integrating 2-photon-derived collagen architecture with vascular mechanical measurements in the murine pulmonary artery. Although the model is trained on chronological age, it functions as a biological age estimator in the sense that its input features are structural and mechanical properties of the tissue itself, not calendar time. A clock trained on chronological age predicts the age at which a given biological state is expected to occur; deviations between predicted and chronological age therefore reflect accelerated or decelerated tissue aging. This distinction is important: chronological age is fixed, while the biological state of the vasculature is modifiable by disease, intervention, or genetic background, making the model a potential tool for detecting divergence between the two.

Our data makes the mechanistic basis for the changes in PA mechanical parameters intuitive. Arterial distensibility is the fractional change in lumen area per unit pressure change and is influenced by the stiffness and geometry of the wall. As collagen fibers uncrimp, crosslink, and increasingly bear load at physiological pressures with advancing age, distensibility decreases measurably. PA circumferential stiffness quantifies this as the modulus at physiological pressures. PWV, computed from the Bramwell-Hill equation, provides a spatially averaged stiffness index across the vessel length and is sensitive to both local and distributed changes in wall composition. These three parameters collectively triangulate the mechanical age of the vessel from three distinct geometric and loading perspectives, providing information that is complements to the microstructure relevant 2-photon model features.

The lung resistance finding is the surprising result of this study and merits further discussion. The gain of +0.194 R² from a single lung mechanics parameter exceeds the gain from any single PA mechanical variable and approaches the total improvement of the three-variable PA mechanical model. Age-related changes in airway resistance reflect progressive ECM remodeling, including peribronchial collagen deposition, increased airspace compliance heterogeneity, and reduced alveolar elastic recoil, driven by the same fibrotic and ECM-remodeling pathways that disorganize RPA collagen [17]. The pulmonary vasculature and airways are anatomically contiguous, share a common interstitial space, and are subject to common aging-associated molecular signals including TGF-β, collagen crosslinking enzymes (LOX family), and matrix metalloproteinase dysregulation [18]. The shared biology of pulmonary vascular, airway, and lung parenchymal aging suggests that biomechanical properties like lung resistance and RPA collagen architecture identify the same underlying tissue remodeling program in separate but complementary compartments of the lung. Distensibility and stiffness reflect the direct mechanical consequence of collagen structural changes in the vessel wall itself, while lung resistance reflects the organ-level functional consequence of pulmonary ECM aging in the airways, a distinct but complementary signal that may explain why lung resistance provides a larger R² gain than distensibility alone despite measuring a different compartment.

The inability to combine PA mechanical and lung mechanics parameters in a single model reflects a technical limitation of this dataset, as they were measured in separate animal cohorts with no overlap, rather than a biological constraint. A prospective study measuring 2-photon microstructure, biaxial mechanics, and flexiVent on the same animals would be expected to yield a combined model with a better LOO R² .The sex dimorphism in the microstructural model (female R² = 0.767 versus male R² = 0.375) is a biologically important finding that survived all multimodal model variants. κ undergoes a nonlinear post-18-month decline in female group that produces high feature contrast, potentially reflecting age-associated alterations in estrogen-mediated regulation in aging female C57BL/6J mice [19]. The absence of a comparably strong signal in males may reflect genuinely slower or later-onset collagen disorganization in the male RPA. A prospective study with matched sex representation and complete mechanical measurements is necessary to definitively characterize whether the sex difference in biological aging rate is real or dataset dependent.

The pattern of model performance across sex and modality carries a specific mechanistic implication that extends beyond a statistical observation. In females, the 2-photon-only model achieves LOO R² = 0.767 (Figure 3C), indicating that collagen fiber orientation and straightness alone encode the majority of age-predictive information in the female RPA. Adding vascular mechanical parameters raises female R² to 0.960, a gain of +0.193 that reflects mechanical confirmation of a structurally relevant aging program. In males, by contrast, the 2-photon-only model achieves only R² = 0.375 (Figure 3B), despite the male animals spanning the same chronological age range; the flat κ trajectory visible in Figure 3B confirms that collagen anisotropy does not reorganize systematically over this lifespan window. Adding mechanical parameters recovers male R² to 0.658, a gain of +0.283 that substantially exceeds the female gain which demonstrates that male vascular aging is mechanically real and temporally ordered but poorly captured by 2-photon collagen imaging. Taken together, these findings indicate that collagen architectural remodeling is the primary aging clock mechanism in the female murine RPA, while the male aging program operates through processes that are captured by bulk mechanical measurements but not by fiber-level structural imaging. This distinction is biologically plausible: estrogen modulates collagen synthesis, LOX-mediated crosslinking, and MMP activity in vascular tissue [19]. The male vascular wall may age equivalently in a functional sense with an example of stiffening on schedule as shown by the mechanical parameters, while the underlying fiber architecture responds more slowly or heterogeneously, reducing the collagen features’ capacity to serve as a reliable age clock in that sex.

Several limitations should be acknowledged. The multimodal model (n = 36) is evaluated on a smaller sample than the microstructure-only model (n = 81), and the R² comparison across configurations with different *n* must be interpreted with caution. The lung mechanics and PA mechanical cohorts cannot be combined due to a lack of sample overlap, precluding evaluation of the most comprehensive possible model. Individual-level mechanical and 2-photon measurements are available for only a subset of animals, because not all measurement modalities were applied to every animal. Finally, the model has not been validated in pathological vascular aging models or in a different mouse strain.

A consistent pattern in the predictions is that the youngest animals are slightly overpredicted, and the oldest animals slightly underpredicted, compressing the predicted age range relative to the true range. This is the expected signature of regression to the mean, which arises in any regularized regression: the ε-insensitive loss and L2 penalty (C-controlled) shrink predictions toward the training-set mean, and because support vector regression with an RBF kernel cannot extrapolate beyond the range of its support vectors, estimates near the extremes of the age distribution are pulled inward. The effect is most visible at the boundaries because those samples have neighbors on only one side, biasing the locally weighted prediction toward the interior. The resulting residuals are small and approximately symmetric (μ = −0.47 mo, σ = 2.65 mo; Shapiro-Wilk p = 0.303), and the boundary bias does not compromise the model’s ability to rank animals by age or to detect deviations from the normative trajectory. In principle, a second-stage model trained on the first-stage residuals could partially correct this shrinkage but given the small magnitude of the residuals relative to the biological variability of the cohort, we judged the added model complexity and attendant overfitting risk to be unwarranted at this sample size.

Future directions include: (1) a prospective study collecting 2-photon imaging, biaxial mechanical testing, and flexiVent lung mechanics on the same animals, which would enable a fully combined multimodal model and is expected to yield R² exceeding 0.90; (2) expanding male cohorts with complete endpoint data to resolve the sex dimorphism question; (3) applying the von Mises SVR framework to additional vascular beds; and (4) testing how predicted biological age diverges from chronological age in models of cardiopulmonary injury or disease.

In conclusion, integrating 2-photon-derived collagen orientation features with vascular mechanical parameters substantially improves biological age prediction in the murine RPA, raising LOO R² from 0.596 (2-photon only) to 0.834 (multimodal). The finding that lung resistance provides the single largest gain (+0.258 R²) reveals a shared aging biology between pulmonary vascular and parenchymal tissues that may prove useful for future multimodal aging biomarker development. The sex dimorphism in collagen-based aging dynamics represents an important biological finding motivating prospective sex-balanced studies.

## Data and Code Availability

Data and Code available at time of peer-review publication.

## Funding Acknowledgements

This work was supported by the VA VISN1 Fred Wright CDA1, National Institute on Aging R03AG074063, and EPM is a Pepper Scholar of the Yale Claude D. Pepper Older Americans Independent Center supported by NIA P30AG021342. This work is supported by NSF grant 2436623 to ABR.

## Author Contributions

P.D.: conceptualization, analysis, modeling, visualization, writing.

JDP: Experiments for validation data, lung data analyses

DL, LL: Lung data analyses

CC: performed experiments for two photon imaging, analyzed data, reviewed and edited manuscript

ABR: performed experiments, edited manuscript

XY: supervision, editing of manuscript

E.P.M.: supervision, funding, review and editing.

*Alt title: Multimodal SVR for murine cardiopulmonary aging*

